## Supplemental Figures for "OsMADS58 stabilizes gene regulatory circuits during rice stamen development"

**SUPPLEMENTAL FIGURES and LEGENDS**

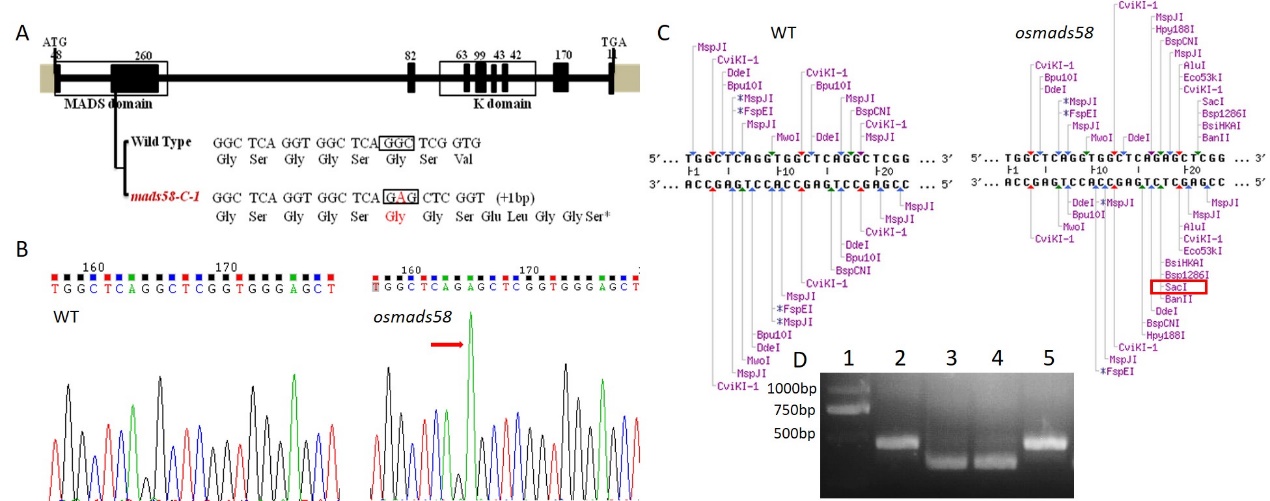

**Supplemental Figure 1.** Null mutation of *OsMADS58* is created by CRISPR technology.

(A) Schematic diagram of the CRISPR constructs designed for *OsMADS58*. Target sequence is located in the second exon. Red “A” represents the inserted nucleotides on the gene in the mutant. The single nucleotide insertion caused a frameshift and a premature translation termination in *osmads58* mutant. (B) The *osmads58* mutation caused by CRISPR were detected using sequencing. The red arrow represents the insertion of a single nucleotide A in the homozygous mutant. (C) The target sequence restriction site display. The homozygous mutant inserted the nucleotide A can be recognized by SacI restriction endonuclease. (D) The *osmads58* homozygous mutant were detected using enzyme digestion. 1. marker; 2 and 5, *OsMADS58* gene seqence in wild-type were digested by SacI; 3 and 4, *OsMADS58* gene seqence in *osmads58* homozygous mutant were digested by SacI.

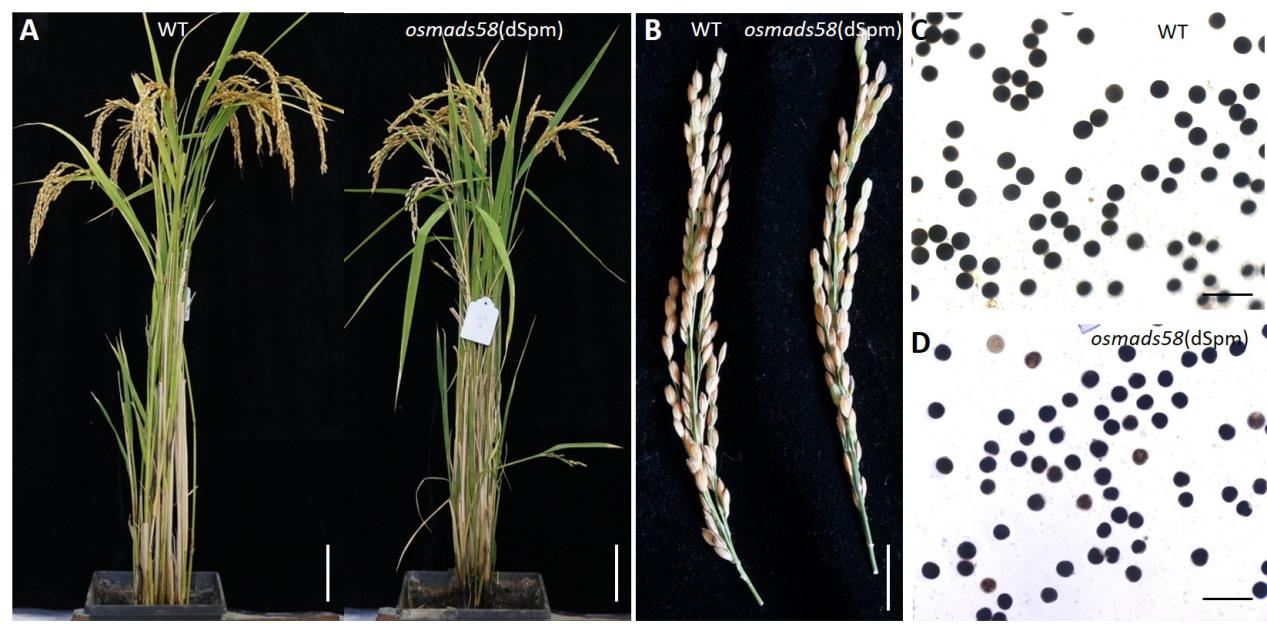

**Supplemental Figure 2.** The *osmads58* dSpm line exhibits normal fertility.

(A) Comparison of wild-type (WT) and *osmads58* dSpm line plants after seed maturation. Bars, 10 cm. (B) Panicles in wild-type and *osmads58* dSpm line plants. Bars, 1cm. (C and D) Wild-type (C) and *osmads58* dSpm line (D) pollen grains were stained by I_2_-KI solution. Bars, 200 μm.

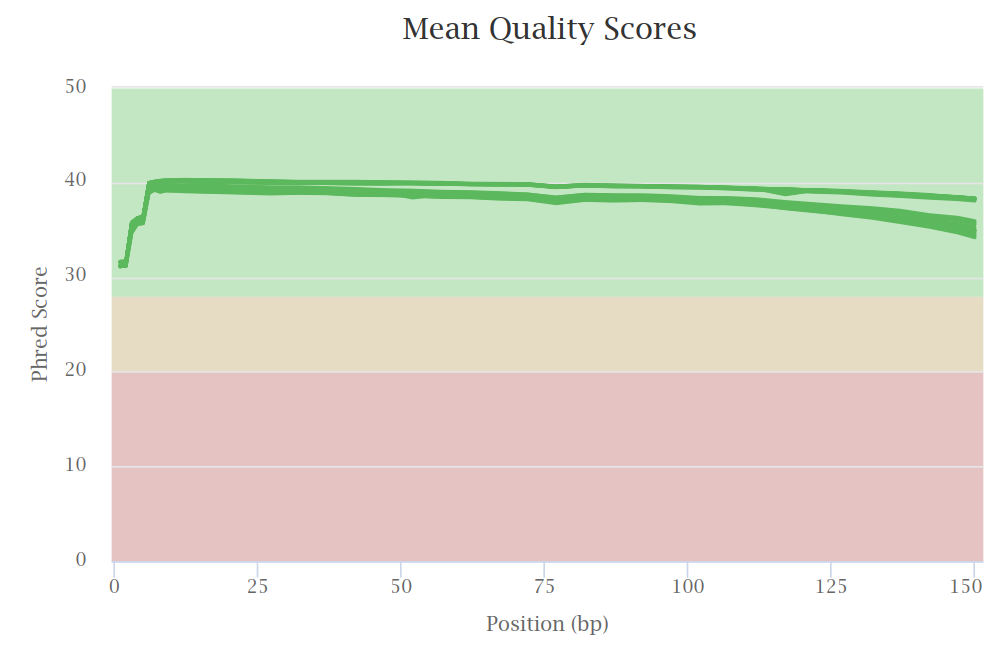

**Supplementary Figure 3.** Average quality (Phred scores) (y-axis) of sequencing reads in each base pair (x-axis).

Each line represents a sample.

­­­­­­­­­­­­­
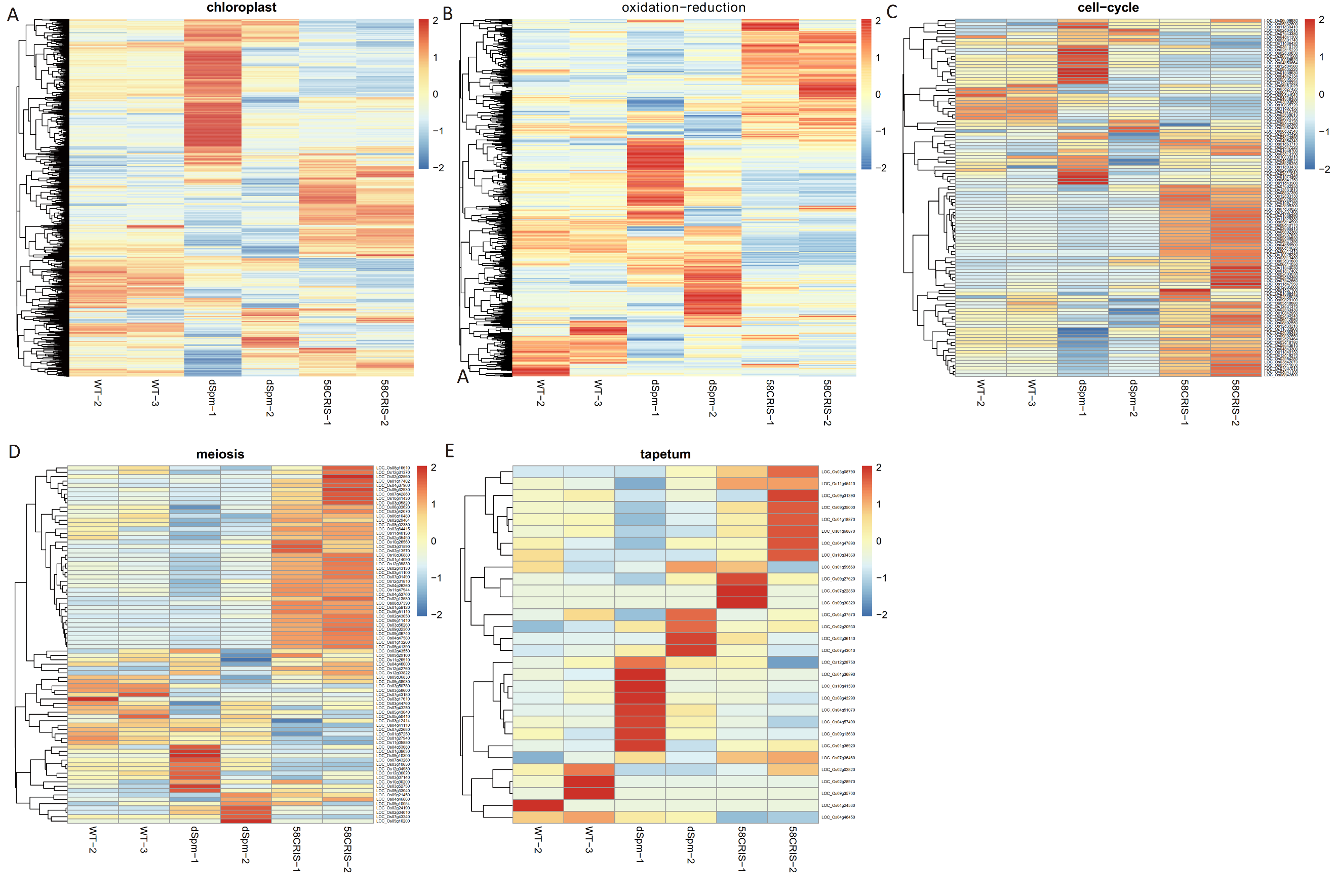

**Supplementary Figure 4.** Expression profiles of genes with known functions in different samples.

The genes related to chloroplast (A), oxidation-reduction (B), cell cycle (C), meiosis (D), and tapetum (E) were shown.
